# Identifying multi-omics biomarkers for ovarian cancer survival estimation

**DOI:** 10.64898/2026.08.04.742866

**Authors:** Kosar Fateh, Srinivasulu Yerukala Sathipati

## Abstract

Ovarian cancer is among the deadliest gynecologic malignancies, and its molecular heterogeneity limits accurate prognostic stratification. Although multi-omics approaches have improved predictive modeling, many prioritize predictive performance over biological interpretability, limiting their clinical translation. We developed an interpretable three-stage machine learning framework integrating mRNA, microRNA, DNA methylation, copy number variation, and protein expression data from The Cancer Genome Atlas. Hierarchical feature selection was combined with a weighted ensemble of ElasticNet, ridge regression, support vector regression, XGBoost, and random forest models to estimate overall survival time in patients with ovarian cancer. Multi-omics integration outperformed every single-modality model, achieving a Pearson correlation of 0.752, a concordance index of 0.779, and a mean absolute error of 8.57 months between estimated and observed survival time, compared with 0.48 for the best single modality. The framework identified a 20-biomarker signature dominated by tumor-associated macrophage and complement genes. In an independent survival analysis, *VSIG4* and *CD163* remained significant after false discovery rate correction, and the signature raised the concordance index over clinical covariates alone from 0.615 to 0.686Enrichment analysis implicated PI3K-Akt, MAPK, focal adhesion, hypoxia, apoptosis, and p53 signaling pathways. This framework couples improved prognostic estimation with biological interpretability supporting multi-omics biomarker discovery in ovarian cancer.

## Introduction

Ovarian cancer remains one of the most lethal gynecologic malignancies worldwide and accounts for a disproportionate number of deaths among female reproductive cancers. Recent global estimates report approximately 325,000 new cases and 235,000 deaths annually, underscoring its substantial clinical burden^1^. Unlike many other malignancies, ovarian cancer is frequently diagnosed only after extensive intraperitoneal dissemination because early-stage disease is typically asymptomatic or presents with vague, non-specific symptoms^2^. Consequently, although patients diagnosed with localized disease have favorable survival outcomes, the majority present with advanced-stage disease characterized by extensive metastasis, frequent recurrence, chemotherapy resistance, and poor long-term survival^2,3^. Improving the ability to identify patients with aggressive disease therefore remains a central challenge in ovarian cancer management.

This clinical heterogeneity reflects the molecular complexity of ovarian cancer, which is driven by coordinated alterations across genomic, epigenomic, transcriptomic, and proteomic layers rather than isolated genetic events. Recurrent alterations in key cancer-associated genes, including *TP53*, *BRCA1*, *BRCA2*, *NF1*, *RB1*, and *CDK12*, disrupt pathways involved in DNA repair, cell-cycle regulation, apoptosis, metabolism, and immune signaling^4^. In addition, interactions within the tumor microenvironment, particularly those involving tumor-associated macrophages, inflammatory signaling, angiogenesis, and extracellular matrix remodeling, further contribute to disease progression and therapeutic response. Together, these interconnected molecular mechanisms suggest that clinically meaningful prognostic biomarkers are unlikely to arise from a single omics layer or isolated biological pathway^5^.

Advances in high-throughput sequencing and proteomic technologies have enabled the generation of large-scale multi-omics datasets, providing unprecedented opportunities to investigate ovarian cancer biology from complementary molecular perspectives^6,7^. Numerous studies have identified candidate biomarkers associated with survival, recurrence, and treatment response^8–10^. However, relatively few have translated into clinical practice because reproducibility remains a persistent challenge. Variability in patient cohorts, specimen processing, analytical methodology, and statistical approach frequently leads to inconsistent findings across independent studies^6^. More importantly, some reported biomarkers remain isolated molecular observations that lack sufficient biological validation and mechanistic context, limiting both their clinical translation and their contribution to understanding disease progression^8,9^.

Integrating multiple omics layers offers a more comprehensive representation of ovarian cancer biology by capturing coordinated alterations across genomic, epigenomic, transcriptomic, and proteomic levels, including copy number variation, DNA methylation, miRNA expression, mRNA expression, and protein abundance^11,12^. However, multi-omics datasets are inherently high-dimensional, heterogeneous and noisy, with molecular features often vastly outnumbering available patient samples. Consequently, robust feature selection and computational modeling are therefore essential to minimize redundancy, reduce overfitting, and identify biologically meaningful biomarkers^11,13^. For instance, El-Manzalawy et al. proposed a multi-view feature selection strategy for ovarian cancer survival prediction that maximized feature relevance while minimizing redundancy in high-dimensional multi-omics data^14^. Recently, Zhou et al. integrated multiple omics modalities with machine learning to identify molecular subtypes and construct prognostic models^15^.

Machine learning has become a powerful tool for analyzing complex multi-omics datasets, enabling the identification of molecular patterns beyond the reach of conventional statistical methods^13,16–17^. Eldesouki et al. integrated clinical and molecular features from TCGA using multiple machine learning algorithms to improve prognostic stratification while identifying clinically relevant biomarkers^18^, and Fu et al. developed integrated multi-omics survival predictors from genomic, transcriptomic, and epigenomic profiles^19^. Representative studies are summarized in **Supplementary Table S1**. Although these studies demonstrate promising predictive performance, many prioritize predictive accuracy over biological interpretability, limiting the clinical translation of identified biomarkers and the mechanistic insight they provide^13,16–17^. Computational frameworks capable of integrating heterogeneous molecular data while identifying robust, interpretable, and biologically meaningful prognostic biomarkers therefore remain needed.

In this study, we developed an interpretable computational framework that integrates five complementary TCGA ovarian cancer omics layers with clinical and survival data to identify robust prognostic biomarkers. A distinguishing characteristic of the framework is that it prioritizes biologically relevant biomarkers, with most selected features having previously reported roles in ovarian cancer progression. Unlike approaches focused solely on prediction, our framework combines multi-omics integration, ensemble machine learning, and survival validation to identify biomarker signatures that are both predictive and biologically informative.

## Results

### Dataset overview, cohort selection and clinical characteristics

The molecular landscape of the TCGA-OV dataset was first characterized across the five omics modalities (**Fig. 1**). The dataset exhibited marked heterogeneity in feature dimensionality, ranging from 131 RPPA proteins to 27,578 DNA methylation probes, while the number of patients available per modality varied from 412 to 592 (**Fig. 1A**). Pairwise correlation analysis based on the first principal component of each modality revealed generally weak inter-modality associations (Pearson *r* = −0.15 to 0.28; **Fig. 1B**), indicating that the different molecular layers capture largely distinct biological information and supporting the integration of complementary modalities.

**Figure 1.**
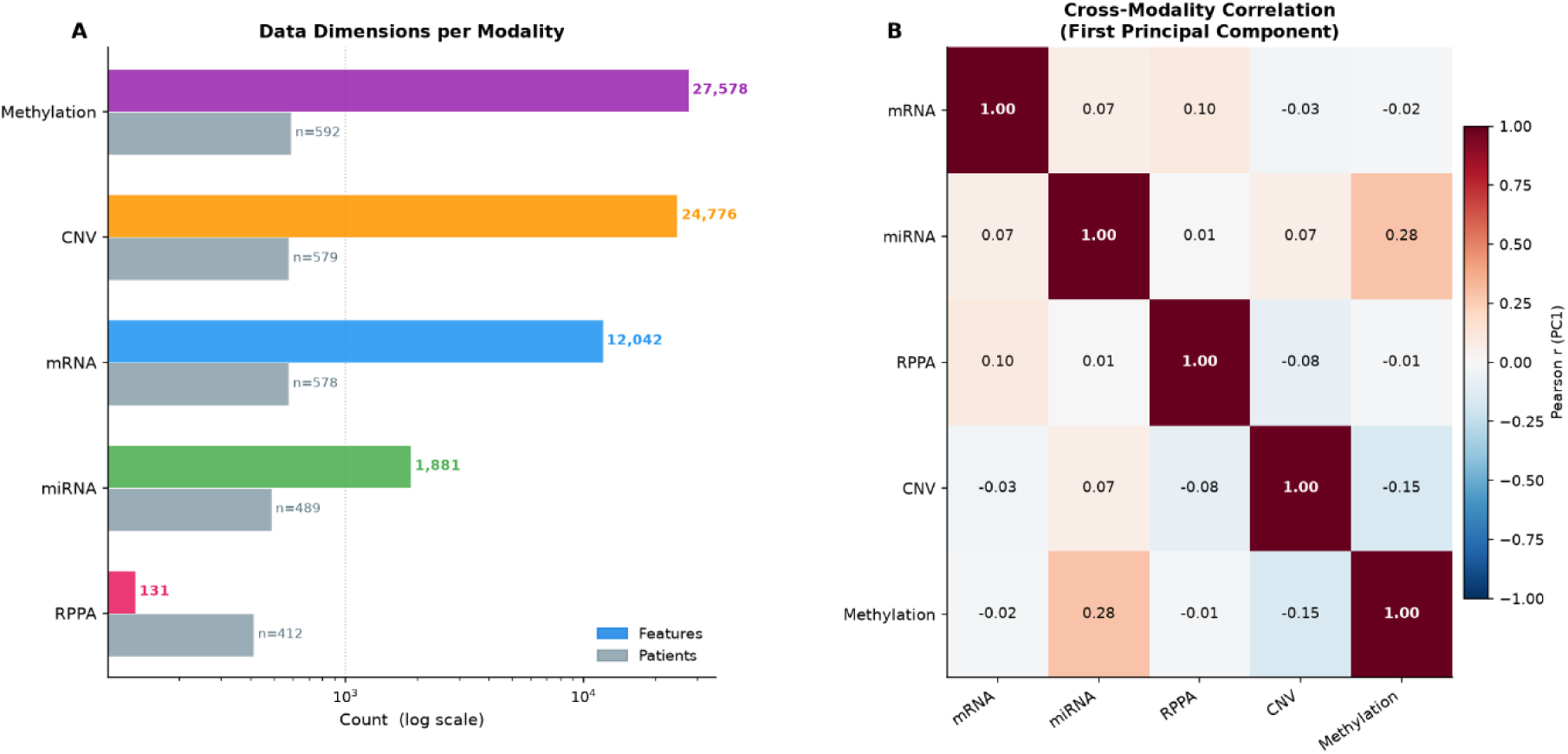
Overview of the TCGA-OV multi-omics dataset. **(A)** Number of molecular features and patient samples available for each omics modality included in this study. The feature counts were 12,042 for mRNA, 1,881 for miRNA, 24,776 for copy number variation (CNV), 27,578 for DNA methylation, and 131 for reverse-phase protein array (RPPA), with corresponding sample sizes ranging from 412 to 592 patients. **(B)** Pairwise Pearson correlation matrix based on the first principal component (PC1) of each omics modality, illustrating the relationships among the five molecular data types.

The TCGA-OV dataset comprised 578 patients with mRNA expression, 489 with miRNA expression, 579 with copy number variation (CNV), 592 with DNA methylation, and 412 with reverse-phase protein array (RPPA) protein expression (**Fig. 2A**). Integrating these datasets yielded a five-omics intersection of 330 patients with complete molecular profiles. After excluding one patient without complete survival information, a 329-patient cohort was established for Kaplan–Meier and Cox proportional hazards analyses, comprising 212 deceased and 117 censored patients (**Fig. 2A**). Clinicopathological characteristics are summarized in **Table 1**.

**Figure 2.**
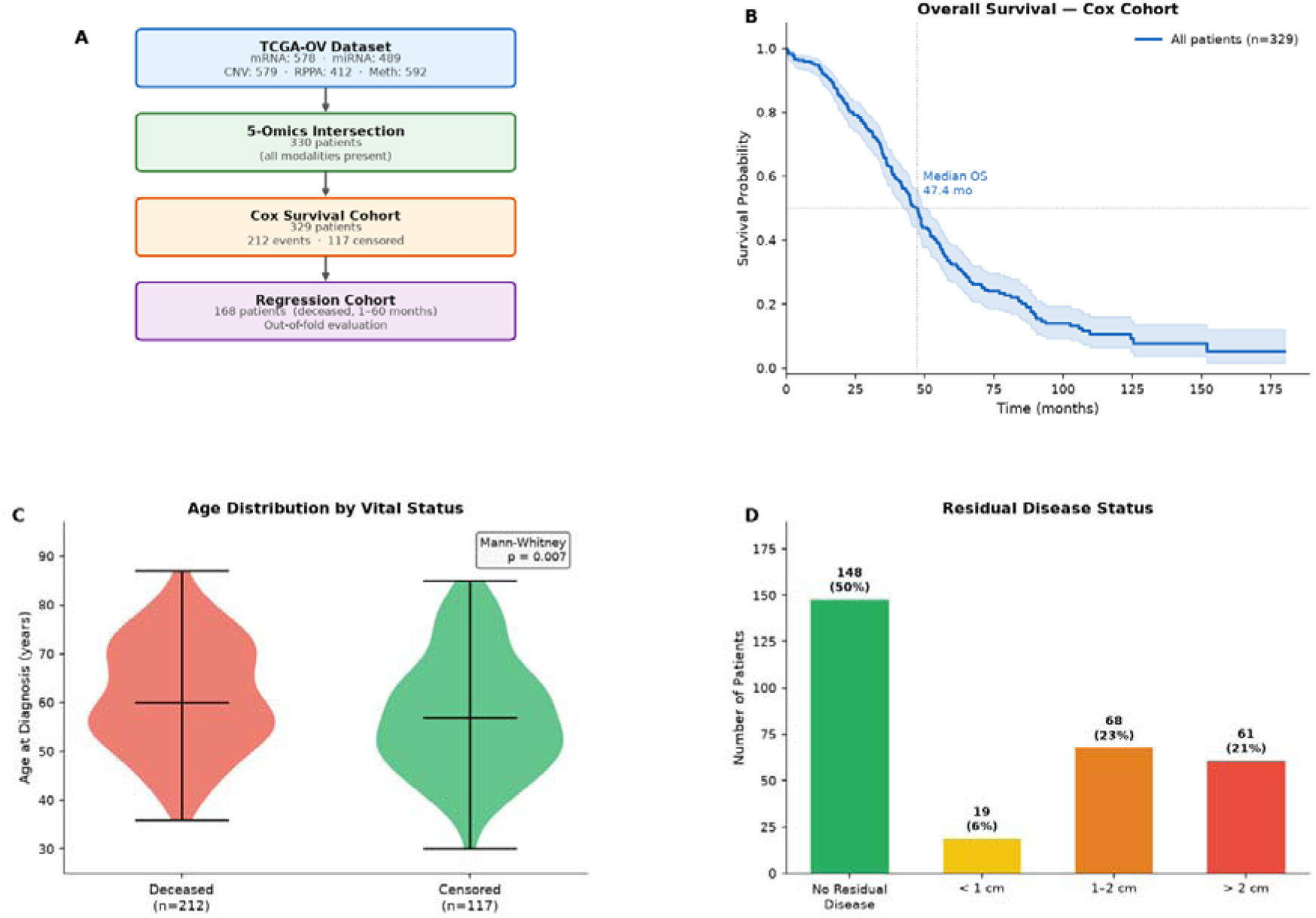
Cohort selection and clinical characteristics of the study population. (A) Workflow illustrating cohort construction from the TCGA-OV multi-omics dataset, including the five-omics intersection cohort, the cohort used for survival analyses, and the final regression cohort used for machine learning model development. (B) Kaplan–Meier overall survival curve of the Cox cohort, with the shaded area representing the 95% confidence interval. (C) Distribution of age at diagnosis according to patient vital status. Statistical significance was assessed using the Mann–Whitney U test. (D) Distribution of residual disease status within the study cohort.

**Table 1.** Clinicopathological characteristics of the 329-patient TCGA ovarian cancer cohort.

| Variable | Category | Value |
| --- | --- | --- |
| **Total patients** |  | n = 329 |
| **Events (deceased)** |  | n = 212 (64.4%) |
| **Censored (alive)** |  | n = 117 (35.6%) |
| **Age at diagnosis (years)** | Mean ± SD | 59.8 ± 11.6 |
| **Age at diagnosis (years)** | Median [IQR] | 59.0 [51.0–69.0] |
| **Overall survival (months)** | Median observed [IQR] | 35.2 [17.4–56.5] |
| **Residual disease** | No residual disease | n = 148 (45.0%) |
| **Residual disease** | < 1 cm | n = 19 (5.8%) |
| **Residual disease** | 1–2 cm | n = 68 (20.7%) |
| **Residual disease** | > 2 cm | n = 61 (18.5%) |
|  | Not available | n = 33 |
| **Tumor grade** | Grade 1 | n = 1 (0.3%) |
| **Tumor grade** | Grade 2 | n = 40 (12.3%) |
| **Tumor grade** | Grade 3 | n = 282 (87.0%) |
| **Tumor grade** | Grade 4 | n = 1 (0.3%) |
|  | Not available | n = 5 |

*IQR, interquartile range; SD, standard deviation. Percentages for residual disease are calculated among the 296 patients with available data, and for tumor grade among the 324 patients with available data.*

Kaplan–Meier analysis of the cohort estimated a median overall survival of 47.4 months (Fig. **2B**); the corresponding raw median of observed follow-up time, which includes censored observations and therefore underestimates survival, was 35.2 months (interquartile range 17.4-56.5). Deceased patients were significantly older at diagnosis than censored patients (Mann–Whitney U test, two-sided, n=329 *p* = 0.007; **Fig. 2C**). Among the 296 patients with available residual disease information, 148 (50.0%) had no residual disease, 19 (6.4%) had residual disease below 1 cm, 68 (23.0%) had residual disease measuring 1–2 cm, and 61 (20.6%) had more than 2 cm (**Fig. 2D**).

Applying the predefined regression criteria, restriction to patients with an observed death event, overall survival between 1 and 60 months, and interquartile-range outlier exclusion, produced a final regression cohort of 168 patients used for machine learning model development and evaluation (**Fig. 2A**). The implications of this outcome-dependent selection are addressed in the Discussion.

### Performance of the multi-omics prediction framework

Predictive performance was assessed for each omics modality and compared with that of the integrated fusion model (**Table 2**). Among single-omics models, mRNA achieved the highest predictive performance (Pearson *r* = 0.481, Spearman ρ = 0.487, C-index = 0.662, mean absolute error [MAE] = 11.09 months) when compared to other modalities. The miRNA, CNV, RPPA, and methylation models performed less well, with methylation weakest (Pearson *r* = 0.050, C-index = 0.528).

**Table 2.** Performance of individual omics models and the multi-omics fusion framework.

| **Model** | **Patients (*n*)** | **Pearson *r*** | **C-index** | **MAE (months)** |  |
| --- | --- | --- | --- | --- | --- |
| **mRNA** | 168 | 0.481 | 0.662 | 11.09 |  |
| **miRNA** | 168 | 0.367 | 0.617 | 11.95 |  |
| **Methylation** | 168 | 0.050 | 0.528 | 12.98 |  |
| **CNV** | 168 | 0.202 | 0.554 | 12.76 |  |
| **RPPA** | 168 | 0.211 | 0.570 | 12.66 |  |
| **Multi-Omics Fusion** | **168** | **0.752** | **0.779** | **8.57** |  |
| *MAE,* | *mean* |  | *absolute* |  | *error* |

Integration of the five molecular modalities through the multi-stage fusion framework substantially improved prognostic estimation relative to all individual omics models, achieving the highest Pearson correlation (0.752), Spearman correlation (0.756) and concordance index (0.779; bootstrap 95% CI 0.746-0.808), and the lowest mean absolute error (8.57 months), explaining 52.1% variance in observed survival, as shown in **Figure 3**. Compared with the best single-omics model (mRNA), integration increased the Pearson correlation by 56.3% (0.481 to 0.752), improved the C-index from 0.662 to 0.779, and reduced estimation error from 11.09 to 8.57 months, a 22.7% reduction. The observed-versus-predicted survival plot demonstrated a strong positive linear relationship between estimated and observed survival times (**Fig. 3A**). Bland–Altman analysis showed minimal systematic bias (0.13 months; 95% limits of agreement −20.5 to +20.8 months), with most predictions falling within the limits of agreement (**Fig. 3B**). The estimated survival time distribution closely approximated the observed distribution, with nearly identical medians (34.2 vs. 34.8 months; **Fig. 3C**), indicating that the model preserved the overall survival characteristics of the study cohort. Time-dependent discrimination was consistent across landmarks, with areas under the receiver operating characteristic curve of 0.90, 0.90, 0.88 and 0.88 at 12, 24, 36 and 48 months, respectively.

**Figure 3.**
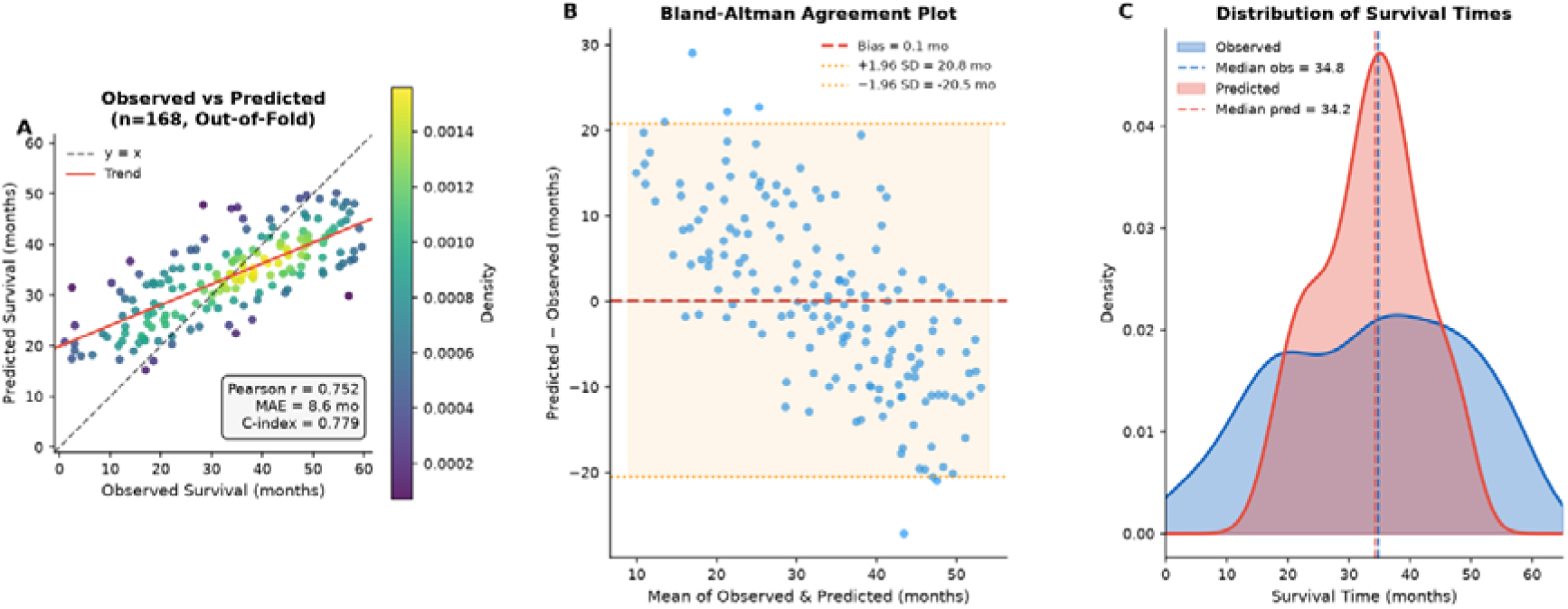
Performance evaluation of the multi-omics survival prediction model. (A) Scatter plot of observed versus out-of-fold predicted overall survival times for the final multi-omics ensemble model (n = 168). The dashed line represents the line of identity (*y* = *x*), and the solid red line indicates the fitted linear trend. Point color represents the local density of observations. (B) Bland–Altman plot showing the agreement between observed and predicted survival times. The red dashed line indicates the mean prediction bias, and the orange dotted lines represent the 95% limits of agreement (±1.96 standard deviations). (C) Kernel density distributions of observed and predicted survival times. Vertical dashed lines indicate the median observed and predicted survival times.

### Prognostic risk stratification

To assess the stratification capacity, the regression cohort was divided into tertiles of estimated survival time. Kaplan–Meier analysis revealed clear and ordered separation among predicted high-, intermediate-, and low-risk groups, with median overall survival of 18.5, 34.8 and 48.1 months respectively (log-rank test across three groups, χ² = 101.8, degrees of freedom = 2, *n* = 168, P = 7.9 × 10^-23^; **Fig. 4**). The high-risk group had the poorest survival outcomes, the low-risk group had the most favorable outcomes, and the intermediate-risk group showed an intermediate survival pattern. Modeled as a continuous variable, each standard deviation increase in predicted risk corresponded to a hazard ratio of 2.91 (95% CI 2.34–3.62, P = 7.4 × 10^-21^). Because this analysis was conducted within the regression cohort, in which all patients experienced the event by construction, it characterizes the internal consistency of the model rather than providing independent prognostic validation.

**Figure 4.**
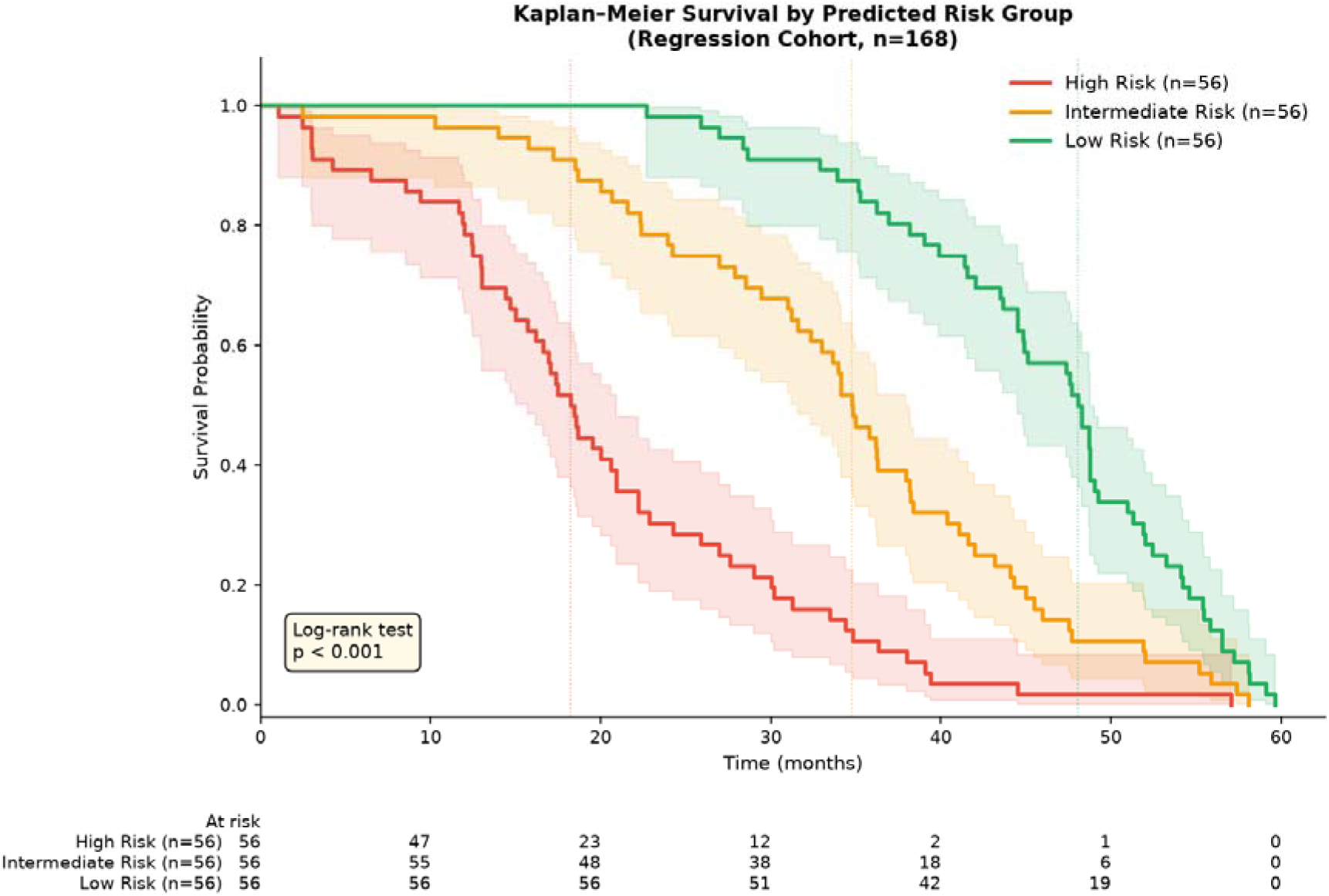
Kaplan–Meier survival curves stratified by predicted risk groups. Patients in the regression cohort (*n* = 168) were classified into high-, intermediate-, and low-risk groups according to the survival times predicted by the multi-omics ensemble model. Kaplan–Meier survival curves were generated for each predicted risk group, with shaded regions representing 95% confidence intervals. The numbers of patients at risk at each time point are shown below the plot. Vertical dotted lines indicate the median survival time for each predicted risk group. Survival differences among the three groups were assessed using the log-rank test.

### Identification of Prognostic Biomarkers

The multi-stage feature selection and ensemble learning framework identified a 20-biomarker signature comprising 13 mRNA and 7 miRNA features, each selected in 100% of cross-validation folds (**Table 3**). The signature was dominated by genes associated with tumor-associated macrophages and complement activation, including *CD163, VSIG4, C1QA, C1QB, TYROBP* and *CD302*.

**Table 3.**
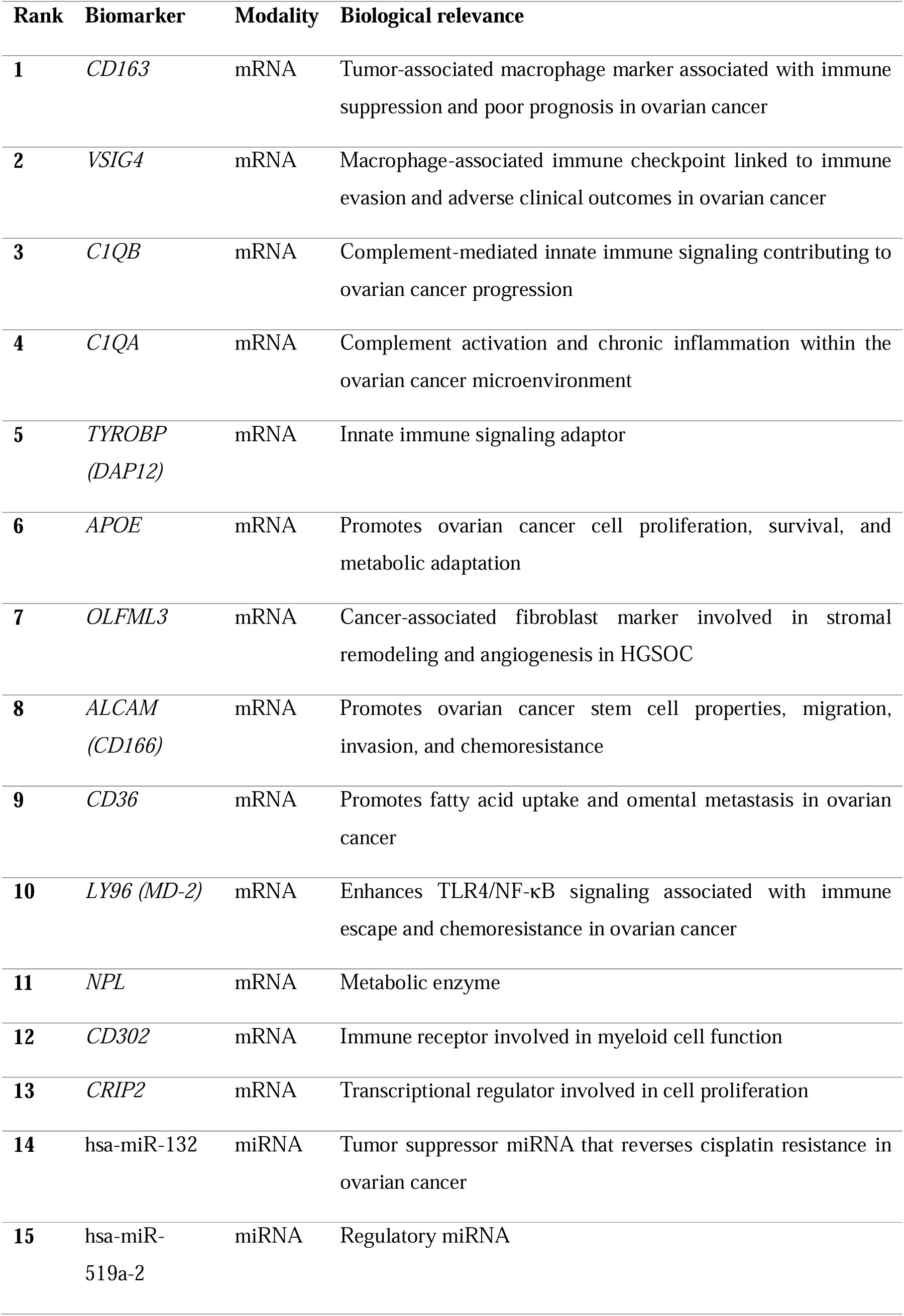
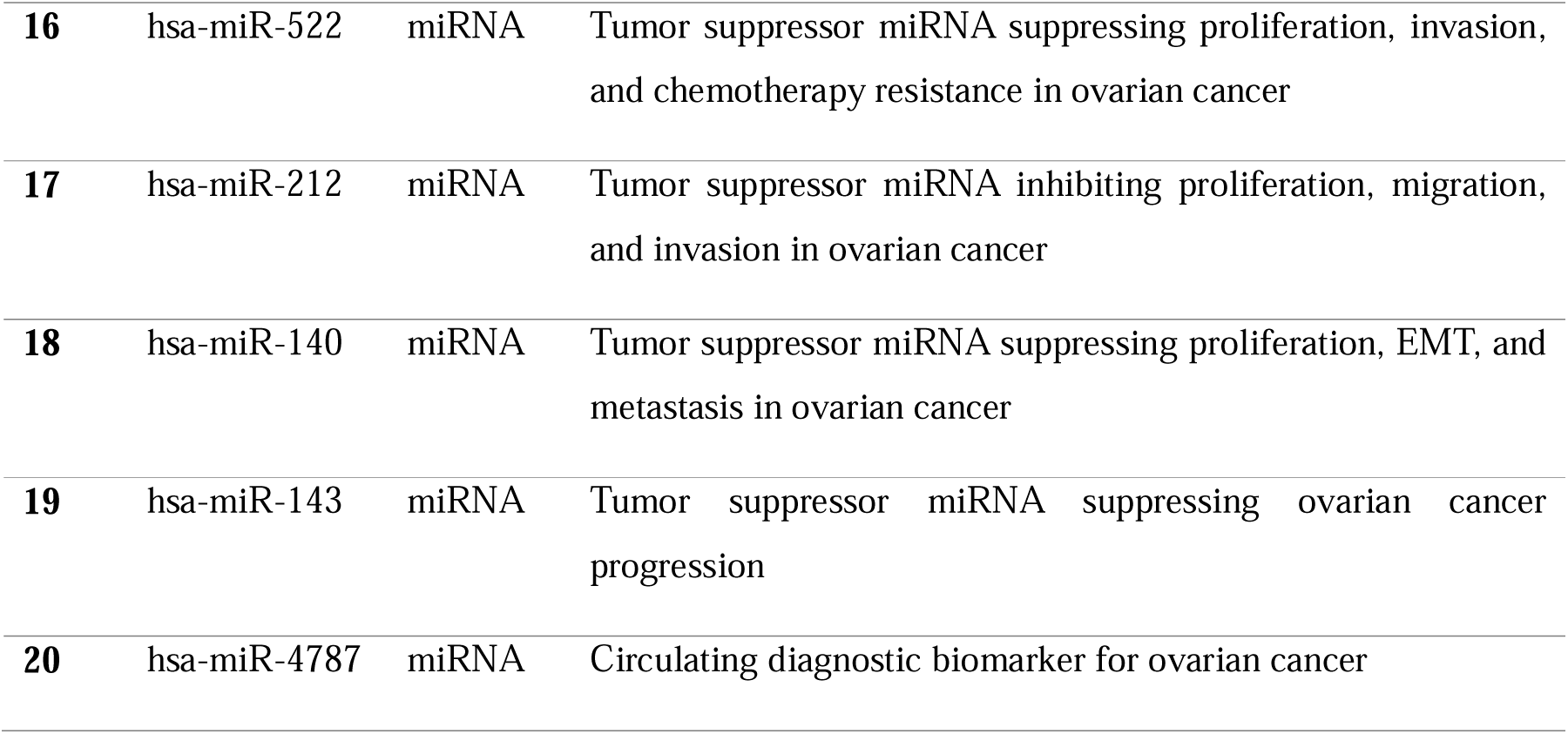
Multi-omics prognostic biomarker signature.

### Survival validation of the signature

To evaluate prognostic significance independently of the regression cohort, each signature biomarker was tested in the full 329-patient cohort, which includes the 117 censored patients and therefore handles incomplete follow-up correctly through the Cox partial likelihood. In univariate Cox proportional hazards models with Benjamini–Hochberg correction across all 22 tested variables, age was the strongest prognostic factor (HR = 1.404 per standard deviation, 95% CI 1.211–1.628, P = 7.0 × 10^-6^, q = 1.5 × 10^-4^). Two signature biomarkers remained significant after correction: *VSIG4* (HR = 1.319, 95% CI 1.139–1.527, P = 2.1 × 10^-4,^ q = 0.002) and *CD163* (HR = 1.238, 95% CI 1.072–1.429, P = 0.004, q = 0.026). Hsa-miR-143 (HR = 1.199, P = 0.015, q = 0.081), C1QB (HR = 1.151, P = 0.046, q = 0.189) and hsa-miR-132 (HR = 1.140, P = 0.052, q = 0.189) showed consistent directional trends that did not survive correction (**Supplementary Table S2**).

In a multivariable L2-penalised Cox model containing all 20 biomarkers together with age and residual disease, age (HR = 1.367, P = 2.1 × 10^-5^), VSIG4 (HR = 1.388, 95% CI 1.080–1.784, P = 0.011) and hsa-miR-143 (HR = 1.199, 95% CI 1.023–1.405, P = 0.025) retained independent associations with overall survival (Supplementary Table S2). Notably, the two lead markers of the signature, *VSIG4* and *CD163*, are the canonical M2 macrophage markers, and their independent replication in the censoring-aware cohort supports the biological coherence of the machine-selected panel. Adding the signature to the clinical covariates increased the concordance index from 0.615 to 0.686, an absolute gain of 0.070, indicating that the molecular features carry prognostic information beyond age and residual disease.

### Correlation of the machine learning-derived transcriptomic biomarkers

The relationship between the transcriptomic biomarkers was performed using the patient-level standardized mean expression values of the selected mRNA and miRNA biomarkers (**Supplementary Fig. S1**). A statistically significant moderate positive correlation was observed between the integrated mRNA and miRNA expression profiles (Pearson’s *r* = 0.433, *p* = 4.47 × 10^-9^). While the significant association demonstrates overall agreement between the two transcriptomic layers, the moderate correlation coefficient indicates that each modality contributes distinct molecular information.

### Expression profiles across survival groups

Expression of the signature biomarkers was compared across three survival groups (1–24, >24–48, and >48–60 months) using the Kruskal–Wallis test followed by Dunn’s post hoc analysis with Benjamini–Hochberg false discovery rate (FDR) correction (**Supplementary Fig. S2A&B**). Of the 20 candidate biomarkers, 11 showed significant differential expression after correction, including *CD163*, *VSIG4*, *C1QA*, *C1QB*, *TYROBP, ALCAM, LY96, NPL, CD302, CRIP2*, and hsa-miR-4787 (q < 0.05). The remaining biomarkers did not differ significantly among survival groups. Post hoc analysis indicated that most significant differences occurred between the shortest-and longest-survival groups (Groups 1 and 3). Several biomarkers, including *CD163, VSIG4, C1QA, C1QB, TYROBP, LY96*, and *NPL*, also differed significantly between the intermediate-and long-survival groups (Groups 2 and 3), whereas *CRIP2* and hsa-miR-4787 primarily distinguished the short-survival group from the other two. The upper group boundary of 60 months is imposed by the regression cohort definition.

### Functional enrichment analysis

Functional enrichment analysis was performed on the integrated gene set comprising the 13 mRNA biomarkers and the predicted target genes of the seven miRNA biomarkers; the miRNA–target gene interaction network is shown in **Supplementary Figure S3**.

Gene Ontology (GO) annotation analysis identified significant enrichment of biological processes relevant to ovarian cancer (**Supplementary Fig. S4**), the most significant being positive regulation of biological process (adjusted *P* = 5.45 × 10^-28^), apoptotic process (adjusted *P* = 4.49 × 10C¹²), regulation of cell population proliferation (adjusted *P* = 1.43 × 10^-11^), cell migration (adjusted *P* = 6.72 × 10^-9^), and response to hypoxia (adjusted *P* = 4.23 × 10^-8^).

Kyoto Encyclopedia of Genes and Genomes (KEGG) pathway analysis identified significant enrichment of cancer-related signaling pathways (**Fig. 5**), most notably pathways in cancer (adjusted *P* = 8.09 × 10^-8^), Focal adhesion (adjusted *P* = 2.74 × 10^-4^), MAPK signaling pathway (adjusted *P* = 2.03 × 10^-3^), PI3K–Akt signaling pathway (adjusted *P* = 3.79 × 10^-3^), and p53 signaling pathway (adjusted *P* = 8.97 × 10^-3^).

**Figure 5.**
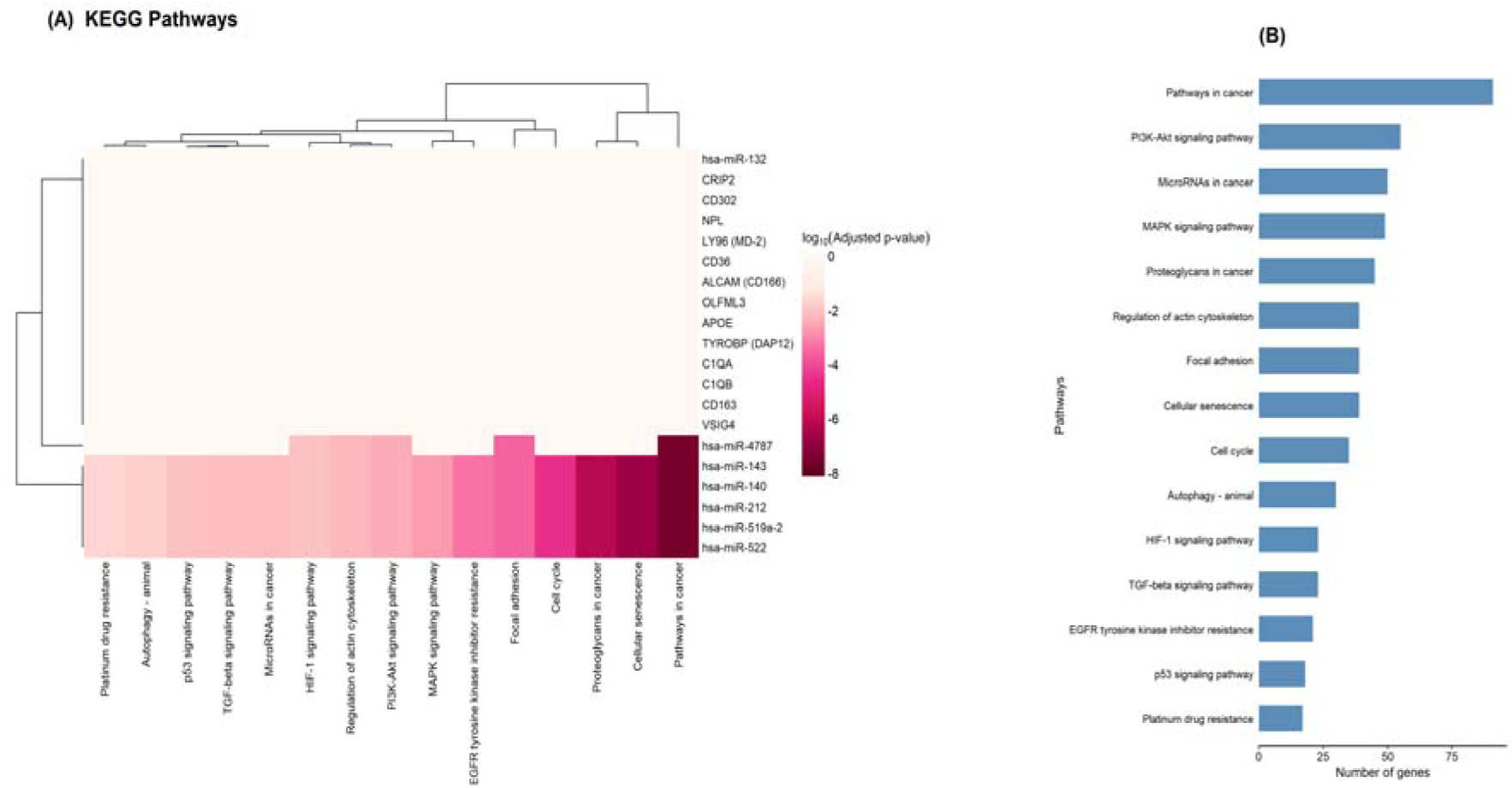
KEGG Pathways Landscape of Candidate Ovarian Cancer Biomarkers. Heatmap (A) showing the associations between candidate mRNA and miRNA biomarkers and the top 15 enriched KEGG pathways. Colored cells indicate biomarker–pathway associations, with color intensity corresponding to the adjusted enrichment *P*-value (darker colors indicate greater statistical significance). Hierarchical clustering was performed on both biomarkers and pathways to identify shared functional patterns. Bar plot (B) summarizes the number of genes contributing to each enriched KEGG pathway.

## Discussion

Accurate prognostic prediction in ovarian cancer remains challenging because clinically relevant molecular information is distributed across multiple regulatory layers rather than concentrated in a single omics modality^13^. In this study, integrating transcriptomic, epigenomic, genomic, and proteomic data substantially improved predictive performance relative to individual omics models. This improvement likely reflects the complementary biological information captured by each layer, and the weak inter-modality correlations we observed support that interpretation of ovarian cancer biology^13^. The staged architecture contributed measurably, allowing a meta-learner to weight modality-level predictions improved correlation beyond simple feature concatenation, suggesting that modality-specific models capture signal that is diluted in a single high-dimensional joint matrix. Similar improvements following multi-omics integration have been reported by Eldesouki et al. and Ren et al., further supporting the value of integrating complementary molecular layers for ovarian cancer prognostic modeling^18,20^.

Rather than yielding a collection of independent markers, our framework revealed a biologically coherent signature in which protein-coding genes and regulatory miRNAs converge on interconnected cellular programs governing ovarian cancer progression. Critically, this coherence was not confined to the cohort in which the signature was derived. When the individual biomarkers were tested in the full 329-patient cohort including censored patients, *VSIG4* and *CD163*, the two canonical markers of M2-polarised macrophages, remained significantly associated with overall survival after false discovery rate correction, and *VSIG4* retained independence in multivariable analysis. That the two strongest survivors of this stricter test are the two markers most specific to the biological program the signature implicates provides a form of internal validation that a purely predictive framework would not supply.

The predominance of immune-related biomarkers, particularly those associated with tumor-associated macrophages and innate immune signaling, is the most notable feature of the signature. Consistent with the observations by Lan et al., Takaishi et al., and Reinartz et al., *CD163*-positive macrophages have been associated with immunosuppressive tumor microenvironments and poor prognosis in ovarian cancer^21–24^. Byun et al. and Wu et al. identified *VSIG4* as a macrophage-associated marker linked to adverse clinical outcomes^25,26^. C1QA and C1QB participate in complement-mediated innate immune signaling linked to chronic inflammation and tumor progression^27,28^. In addition, *LY96* (MD-2), the co-receptor of *TLR4*, may amplify NF-κB-mediated inflammatory signaling and contribute to immune escape and chemoresistance^29,30^. These findings suggest that immune dysregulation, rather than any isolated immune markers, represents a central biological component of the identified prognostic signature^28^.

Several biomarkers associated with tumor dissemination and metastatic progression. *ALCAM* regulates cancer stem cell properties, including self-renewal, adhesion, migration, tumorigenicity, and chemoresistance through activation of the FAK/SRC/paxillin signaling axis^31–34^, whereas Govindarajan et al. demonstrated that *OLFML3* is enriched in cancer-associated fibroblasts in high-grade serous ovarian cancer (HGSOC), supporting a role in stromal remodeling and angiogenesis^35^. Ladányi et al. showed that adipocyte-induced *CD36* expression promotes ovarian cancer metastasis through metabolic reprogramming within the omental microenvironment^36^. *APOE*, likewise, contributes to ovarian cancer cell proliferation, survival, and metabolic adaptation and has been associated with tumor progression and poor prognosis across multiple malignancies^37–39^. Together, these findings emphasize that progression is critically influenced by reciprocal interactions with the stromal and metabolic components of the tumor microenvironment^34–36^. Such complex biological interactions are further refined through post-transcriptional regulation.

The identified miRNA component reinforces this concept. Except for hsa-miR-4787-3p, proposed as a circulating diagnostic biomarker^40^, the remaining miRNAs, including hsa-miR-132, hsa-miR-212, hsa-miR-140, hsa-miR-143, and hsa-miR-522-3p, have been experimentally validated as tumor suppressors in ovarian cancer^41–49^, regulating proliferation, invasion, epithelial–mesenchymal transition (EMT), cell-cycle progression, and chemotherapy resistance through distinct downstream targets^41–46,48,49^. In particular, Zhang et al. demonstrated that hsa-miR-132 reverses cisplatin resistance by targeting *BMI1*, and Shi et al. reported that hsa-miR-143 suppresses ovarian cancer progression^41,49^. It is notable that hsa-miR-143 was among the few features to retain significance in multivariable Cox analysis, aligning the statistical and mechanistic evidence.

Functional enrichment confirmed convergence on shared biological processes and signaling pathways, demonstrating that the biomarkers participate in coordinated oncogenic programs rather than isolated biological functions^50,51^. Enrichment of positive regulation of biological processes, regulation of cell population proliferation, and Pathways in cancer indicates that the signature reflects integrated mechanisms governing sustained proliferation, cell survival, angiogenesis, immune evasion, metabolic adaptation, invasion, and metastatic dissemination^50–52^. Our findings are consistent with recent pathway analyses by Lee et al. and the comprehensive review by Lliberos et al., both of which identified and stated PI3K/Akt and MAPK as central regulators of ovarian cancer progression and therapeutic resistance^51,52^. Specifically, the PI3K/Akt pathway promotes tumor cell survival, proliferation, metabolic adaptation, angiogenesis, and resistance to apoptosis through enhanced DNA repair and pro-survival signaling^50,51,53,54^ whereas the MAPK pathway regulates proliferation, EMT, migration, differentiation, and therapeutic resistance^50–52,55^. Crosstalk between these pathways provides a mechanistic explanation for their simultaneous enrichment in both GO and KEGG analyses^51,52,54^. Beyond promoting proliferation, these pathways also regulate tumor dissemination through their effects on cell adhesion and migration^50–52^.

Ovarian cancer progression is further driven by the coordinated interplay between hypoxia, cell migration, and focal adhesion, which collectively promote tumor invasion and metastatic dissemination. Hypoxia promotes angiogenesis, metabolic adaptation, immune evasion, and therapeutic resistance through activation of hypoxia-inducible factors (HIFs), while also enhancing invasive behavior^53,56^. These effects are mediated in part by PI3K/Akt and MAPK signaling, which coordinate cell motility, proliferation, and survival^52^. Consistent with this, enrichment of the Focal adhesion pathway highlights the critical role of integrin-and focal adhesion kinase (FAK)-mediated interactions with the extracellular matrix in regulating cell adhesion, migration, invasion, and metastatic dissemination, making focal adhesion signaling an attractive therapeutic target in ovarian cancer^50,51^. Finally, enrichment of the apoptotic process and the p53 signaling pathway highlights the central role of defective genomic surveillance and apoptosis evasion in ovarian cancer progression^50,51,57^. Recently, Ma et al. demonstrated that HIF-2α-mediated *TGFBI* activation promotes chemoresistance through PI3K/Akt signaling by simultaneously enhancing DNA repair and suppressing apoptosis^53^. Similarly, Zhang et al. highlighted TP53 dysfunction as a defining molecular event in HGSOC that disrupts genomic surveillance and apoptosis^57^. The prognostic performance of our multi-omics model suggests that they do not arise from a single dysregulated pathway but from coordinated alterations in immune regulation, tumor–microenvironment interactions, post-transcriptional regulation, survival signaling, and genomic stability, which together shape ovarian cancer progression and clinical outcomes^50–52^.

## Limitations

Several limitations should be emphasized. Most importantly, the regression cohort was defined by conditioning on the outcome: only patients with an observed death event, survival between 1 and 60 months, and non-outlying survival times were retained. This design was adopted because the regression target is an observed, uncensored survival time, but it has an important consequence. The reported regression metrics (*r* = 0.752, C-index = 0.779, MAE = 8.57 months) are conditional on the patient having died within five years, a condition that cannot be known at the time a prognostic model would be applied. These values therefore quantify how well the model orders survival times among patients known to have died, and are not directly comparable with censoring-aware concordance indices reported elsewhere, nor should they be read as expected performance in an unselected clinical population. For this reason, we regard the survival analyses in the full 329-patient cohort, where the incremental concordance gain over clinical covariates was a more modest 0.070, as the more clinically meaningful estimate of prognostic value. Future work should fit censoring-aware survival models, such as regularized Cox or random survival forests, to the complete cohort.

Second, the analyses rely exclusively on retrospective TCGA-OV data and require external validation in independent, multi-center cohorts; the mRNA component of the signature is testable in public high-grade serous cohorts. Third, the DNA methylation data derive from the 27K array platform, whose limited genomic coverage may partly explain the weak performance of that modality and should not be taken as evidence that methylation lacks prognostic information. Fourth, hazard ratios from the penalized multivariable model are shrunk toward the null and their confidence intervals are not adjusted for the penalty, so that model is best interpreted as a check on directional consistency rather than a source of precise effect estimates; the univariate results in **Supplementary Table S2** are unpenalized. Finally, the biomarkers were validated computationally and require experimental and prospective clinical confirmation.

## Conclusion

We developed an interpretable multi-stage machine learning framework integrating five complementary omics modalities to identify biologically meaningful prognostic biomarkers for ovarian cancer. The proposed framework substantially improved survival prediction over single-modality models and identified a signature converging on immune regulation, tumor–microenvironment interactions, survival signaling, metastasis, and genomic stability. Two of its lead components, *VSIG4* and *CD163*, replicated as prognostic factors in the full cohort including censored patients, and the signature added measurable discriminative value over clinical covariates alone. This work provides an interpretable framework for multi-omics biomarker discovery that may support prognostic stratification and future precision oncology research in ovarian cancer.

## Materials and Methods

### Data Acquisition

Multi-omics and clinical data for ovarian serous cystadenocarcinoma (TCGA-OV) were obtained from The Cancer Genome Atlas (TCGA) through the UCSC Xena data portal. Five molecular modalities were included in this study: mRNA expression, miRNA expression, DNA methylation (HumanMethylation27), copy number variation (CNV), and reverse-phase protein array (RPPA) protein expression. Corresponding clinical information, including overall survival time and vital status, was also retrieved for all available patients. The complete multi-omics cohort was subsequently used for survival analyses, including Kaplan–Meier survival estimation and Cox proportional hazards regression.

This study used publicly available, de-identified data from TCGA. No new human participants were recruited, and no identifiable personal information was accessed; institutional review board approval and informed consent were therefore not required. All analyses were performed in accordance with the TCGA data use policies and with relevant guidelines and regulations.

### Cohort Selection

Molecular and clinical datasets were harmonized using 12-character TCGA patient identifiers. Only patients with complete molecular profiles across mRNA expression, microRNA (miRNA) expression, DNA methylation, copy number variation, and RPPA protein expression were retained, yielding 330 patients; one lacked valid survival data, giving a 329-patient multi-omics cohort used for Kaplan–Meier and Cox analyses.

For machine learning regression, a separate cohort was constructed by retaining only patients with an observed death event (event = 1), restricting overall survival to 1–60 months, and excluding survival outliers using the interquartile range (IQR) method. This produced a final regression cohort of 168 patients. Overall survival was calculated as days from diagnosis to death divided by 30.44.

### Data preprocessing and multi-stage feature selection

All preprocessing and feature selection were performed independently within each cross-validation training fold, using only training patients, to prevent information leakage into held-out data. Within each fold, features with more than 20% missing values were excluded.

Remaining features were ranked by median absolute deviation and the most variable retained per modality (mRNA 4,000; miRNA 500; DNA methylation 4,000; CNV 4,000; RPPA all 131). mRNA values were log-transformed as log1*p*(*max*(*x*,0)); other modalities were used on their native scales (GISTIC2 log-ratios, beta values, and normalized intensities are already appropriately scaled). Missing values were imputed with training-fold medians and features standardized using training-fold means and standard deviations. Clinical covariates (age, tumor grade, residual disease) were identified by name and protected from the log transformation and variance filter; omics columns were prefixed with their modality label to prevent name collisions.

Feature importance was evaluated using a composite ranking score integrating three complementary measures of association with overall survival: Pearson correlation (linear association), Spearman rank correlation (monotonic association), and mutual information (nonlinear dependency). The composite score was calculated as:

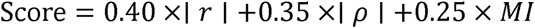

where *r* denotes the Pearson correlation coefficient, _ρ_ denotes the Spearman rank correlation coefficient, and *MI* represents mutual information. Based on this score, the top-ranked features were selected from each modality (50 mRNA, 40 miRNA, 50 DNA methylation, 40 CNV, and 40 RPPA features), and the 25 highest-ranked per modality were carried into the fusion layer. The value K = 25 was chosen by a sweep over K ∈ {15, 20, 25, 30} evaluated by out-of-fold correlation.

### Ensemble Machine Learning Framework

An ensemble of five supervised regression algorithms was used to capture complementary linear and non-linear relationships. ElasticNetCV was incorporated because it combines L1 (LASSO) and L2 (Ridge) regularization with cross-validated hyperparameter optimization, providing effective variable selection while maintaining stability in high-dimensional datasets with correlated predictors^58^It contributed 30% of the ensemble prediction. Ridge regression (α = 1.0) applies L2 regularization to reduce estimator variance under multicollinearity datasets^58, 59^ and contributed 15%.

Support vector regression with a radial basis function kernel (C = 2.0, ε = 1.0, γ = scale) captures non-linear relationships through implicit mapping to a higher-dimensional space ^60,61^, and contributed 25%. XGBoost (500 trees, learning rate 0.02, maximum depth 2, λ = 3.0) sequentially constructs regularized decision trees and contributed 20%^62^.

Random forest regression (300 trees, maximum depth 4) reduces variance through bootstrap aggregation ^63,64^, and contributed the remaining 10%. The prediction is the weighted average of the five components. Ensemble weights were fixed a priori and not tuned on the outcome. Replacing the two linear components with gradient-boosted trees reduced out-of-fold correlation from 0.720 to 0.587, confirming that the dominant signal is approximately additive in log-expression space.

### Three-stage fusion architecture

The framework operates in three stages within a single cross-validation loop that shares fold assignments across all modalities and stages, so that out-of-fold predictions at every level are generated on identical held-out patients. In Stage 1, each of the five modalities is modelled independently as described above, producing five out-of-fold prediction vectors covering all 168 patients. In Stage 2 (early fusion), the top 25 features per modality are concatenated with the clinical covariates and with the five Stage-1 out-of-fold predictions as additional columns, and the same ensemble is trained on this joint matrix. In Stage 3 (late fusion), a meta-learner using the same five-component ensemble is trained on a stacked matrix of six columns: the five Stage-1 out-of-fold predictions and the Stage-2 out-of-fold prediction. The Stage-3 output is the final patient-level prediction. Reporting both stages separately isolates the contribution of stacking.

### Cross-validation and model evaluation

Ten-fold cross-validation was used throughout. Because random K-fold assignment can produce folds with markedly different survival distributions, folds were assigned deterministically: patients were sorted by observed survival time and assigned by round robin, so that patient of rank *i* was placed in fold *i* mod 10. This ensures each fold contains comparable proportions of short-, medium-and long-term survivors and requires no random seed. Preprocessing, feature selection, model training and prediction were performed independently within each training fold. Performance was evaluated using the Pearson correlation coefficient, Spearman rank correlation, concordance index and mean absolute error, computed on out-of-fold predictions. Bootstrap 95% confidence intervals for the concordance index used 2,000 resamples with replacement (seed 42), with bounds at the 2.5th and 97.5th percentiles. Time-dependent discrimination was assessed at 12, 24, 36 and 48 months by treating death at or before each landmark as the positive class and scoring by negative predicted survival.

### Survival Analysis

Survival analyses were performed in the 329-patient cohort, which retains the 117 censored patients and handles incomplete follow-up through the Cox partial likelihood. All covariates were z-score standardized so hazard ratios are expressed per standard deviation and are comparable across modalities with different measurement scales. Each of the 20 signature biomarkers, together with age and residual disease, was first assessed in a separate unpenalized univariate Cox proportional hazards model; P values across the 22 tests were adjusted using the Benjamini–Hochberg false discovery rate procedure. A multivariable Cox model containing all 22 covariates was then fitted with an L2 penalty of 0.1 to improve numerical stability given the number of covariates relative to events; because penalization shrinks coefficients and the standard errors are not adjusted for the penalty, this model is reported as a check on directional consistency rather than as a source of precise effect estimates. Tied event times were handled by the Efron approximation. Incremental prognostic value was quantified by comparing the concordance index of a clinical-only model (age and residual disease) with that of the clinical-plus-signature model. Kaplan–Meier curves were generated by tertile stratification of predicted survival, with differences assessed by the multivariate log-rank test.

### Biomarker Expression Analysis Across Survival Groups

Patients in the regression cohort were stratified into three survival groups (1–24, >24–48 and >48–60 months). Expression of the signature biomarkers was compared across groups using the Kruskal–Wallis test, with pairwise comparisons by Dunn’s post hoc test and Benjamini–Hochberg correction. Non-parametric tests were used because biomarker expression distributions were skewed and did not satisfy normality assumptions, assessed by inspection of quantile–quantile plots and the Shapiro–Wilk test. Functional enrichment was performed on the 13 mRNA biomarkers together with the predicted target genes of the 7 miRNAs, retrieved using the miRNet platform. Gene Ontology (Biological Process, Molecular Function, Cellular Component) and KEGG pathway enrichment were conducted using g:Profiler, with significance determined by the multiple-testing correction implemented in that platform.

### Functional Enrichment Analysis

To investigate the biological functions and molecular pathways associated with the identified biomarkers, functional enrichment analysis was performed on the final panel of 13 mRNA biomarkers together with the predicted target genes of the seven identified miRNAs. The target genes were retrieved using the miRNet platform, and a miRNA–target gene interaction network was generated to visualize the relationships between the selected miRNAs and their predicted targets. The identified target genes were subsequently integrated with the mRNA biomarker panel to generate a unified gene list for downstream Gene Ontology (GO) and Kyoto Encyclopedia of Genes and Genomes (KEGG) enrichment analyses.

Functional enrichment analysis was conducted using g:Profiler. GO enrichment analysis was performed across the three GO domains, including Biological Process (BP), Molecular Function (MF), and Cellular Component (CC), while KEGG pathway enrichment analysis was used to identify significantly enriched signaling pathways. Statistical significance was determined using the multiple-testing correction implemented in g:Profiler, and significantly enriched GO terms and KEGG pathways were retained for downstream biological interpretation.

### Statistical Analysis

Analyses were performed in Python using pandas, NumPy, SciPy, scikit-learn, XGBoost, statsmodels and lifelines. Unless otherwise stated, tests were two-sided and a P value below 0.05 was considered significant; exact P values are reported throughout, and significance in figures is denoted P < 0.05 (*), P < 0.01 (**) and P < 0.001 (***). Where multiple comparisons were made, the Benjamini–Hochberg false discovery rate was applied and adjusted values are reported as q. Random seeds were fixed at 42 for XGBoost, random forest and bootstrap resampling; fold assignment is deterministic and requires no seed. No large language model was used to generate, analyze, or interpret the data reported in this study.

## Acknowledgments

The authors gratefully acknowledge The Cancer Genome Atlas (TCGA) Research Network for generating and making the multi-omics datasets used in this study publicly available through the UCSC Xena platform. The authors also thank the Marshfield Clinic Research Institute for providing the research environment and computational resources that supported this work.

## Author contributions

K.F. collected and curated the data, implemented the computational framework, performed the statistical and bioinformatics analyses together with S.Y.S., interpreted the results, and wrote the manuscript. S.Y.S. conceived and developed the computational framework, supervised the study, provided methodological guidance, and contributed to data interpretation. Both authors reviewed and approved the final manuscript.

## Data Availability

The datasets analyzed during the current study are publicly available from The Cancer Genome Atlas (TCGA) through the UCSC Xena data portal (https://xenabrowser.net/).

## Funding

This work was supported in part by the Marshfield Clinic Research Institute, Marshfield, WI to S.Y.S.

## Competing interests

The authors declare no competing interests.

## Notes

### Competing Interest Statement

The authors have declared no competing interest.

