## Supplementary Figures S1-S4 and Supplementary Tables S1 and S2 for "Identifying multi-omics biomarkers for ovarian cancer survival estimation"

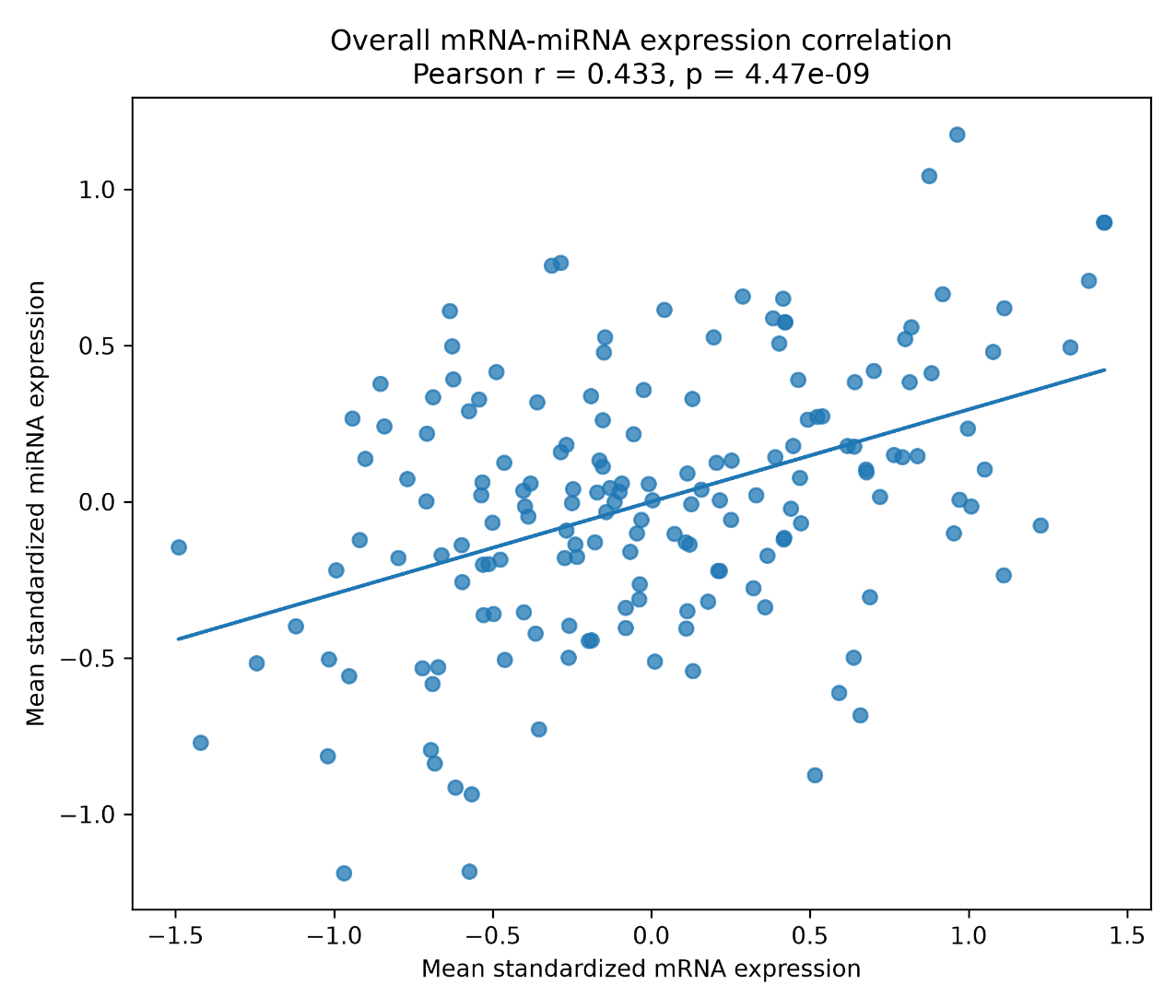
**Supplementary Figure S1. Correlation between the integrated mRNA and miRNA expression profiles.** Scatter plot showing the relationship between the patient-level standardized mean expression values of the selected mRNA and miRNA biomarkers across the study cohort. Each point represents one patient, and the solid line indicates the least-squares linear regression fit. Pearson correlation analysis demonstrated a statistically significant moderate positive association between the two transcriptomic modalities (r = 0.433, p = 4.47 × 10^-9^).

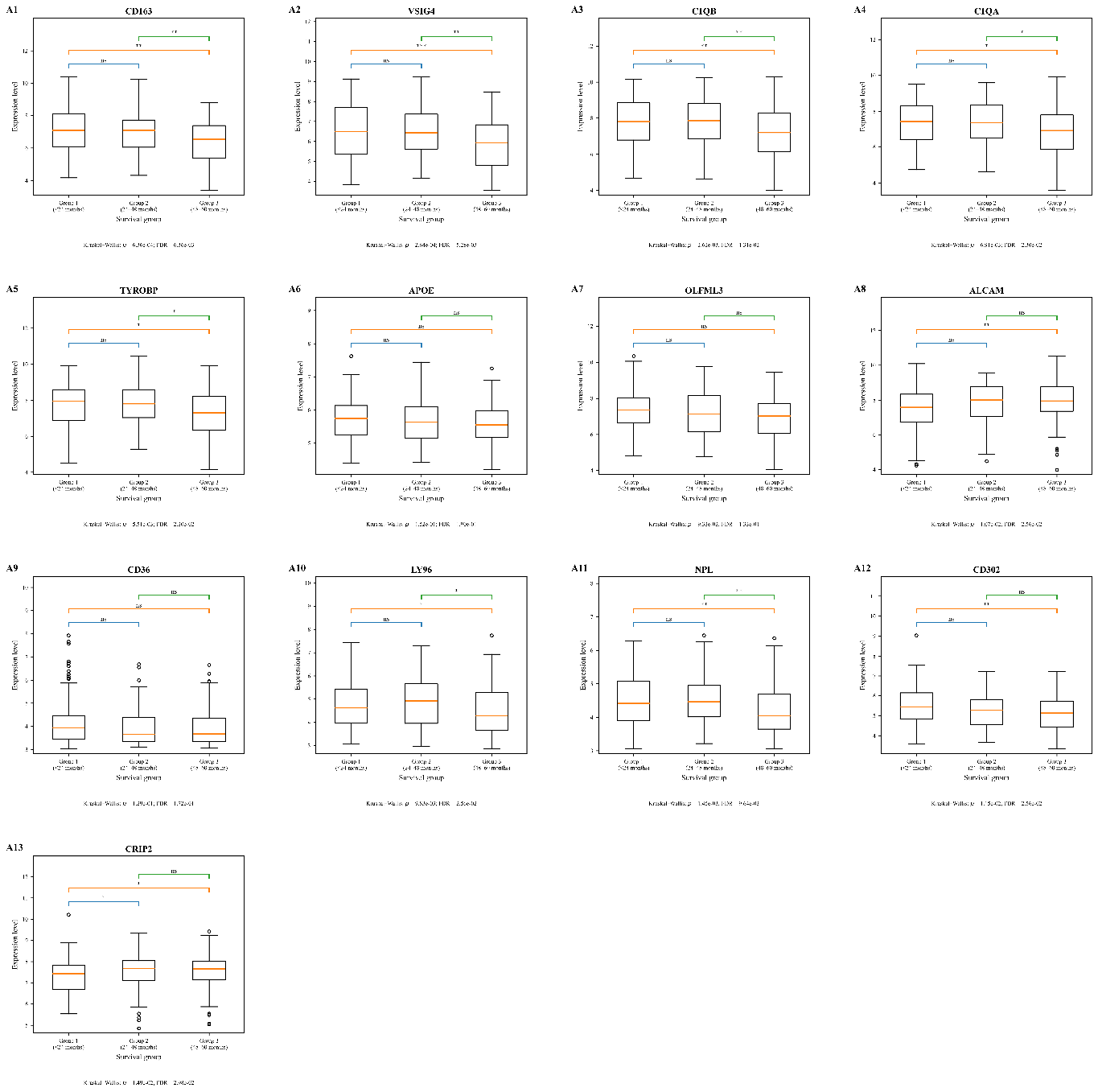
**Supplementary Figure S2A. mRNA biomarker expression across survival groups.**

Boxplots show the expression distributions of the machine learning-selected mRNA biomarkers (A1–A13) among patients with short survival (Group 1, ≤24 months), intermediate survival (Group 2, 24–48 months), and long survival (Group 3, 48–60 months). Differences among the three groups were evaluated using the Kruskal–Wallis test, with Benjamini–Hochberg false discovery rate (FDR) correction across biomarkers. Pairwise group comparisons were performed using Dunn’s post hoc test with FDR adjustment. Center lines indicate medians, boxes represent the interquartile range, whiskers extend to 1.5 times the interquartile range, and circles denote outliers. Pairwise significance is indicated as *p* < 0.05 (**), p < 0.01 (**), p < 0.001 (****), and not significant (ns).

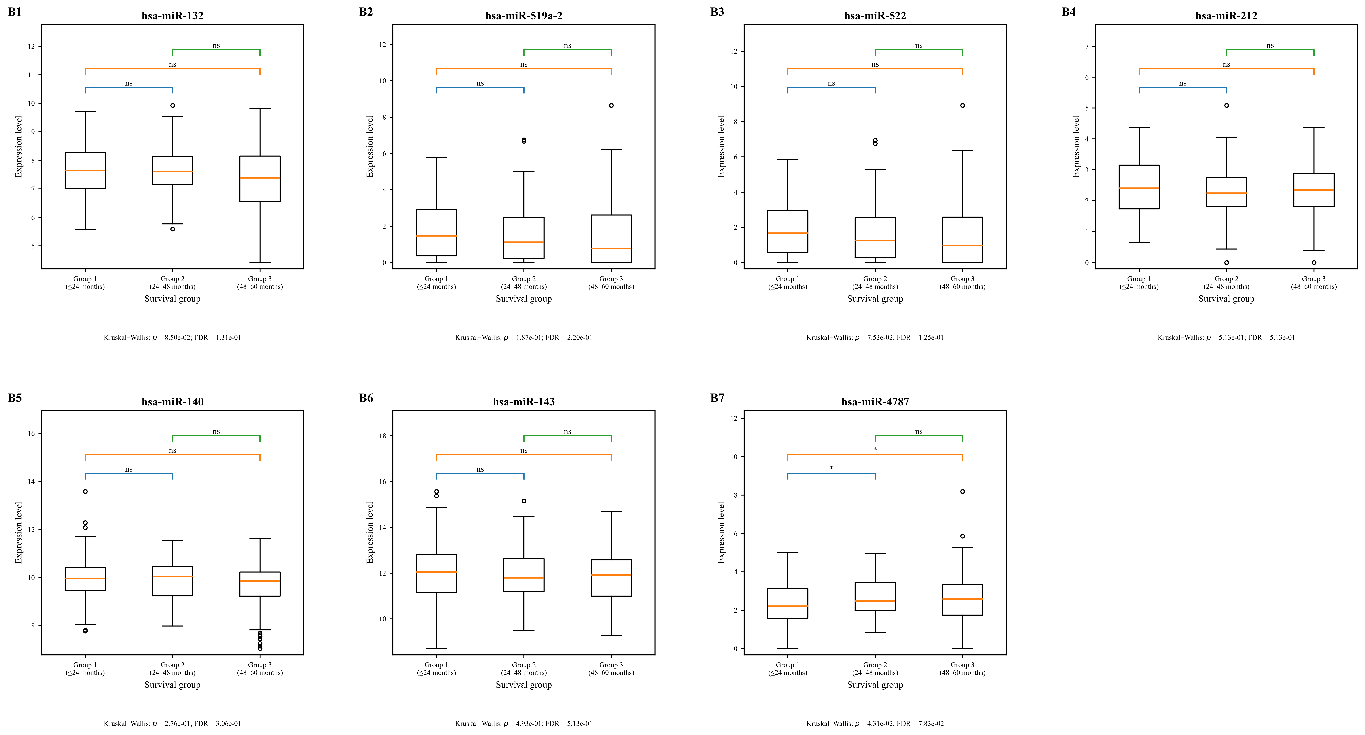
**Supplementary Figure S2B. miRNA biomarker expression across survival groups.** Boxplots show the expression distributions of the machine learning-selected miRNA biomarkers (B1–B7) among patients with short survival (Group 1, ≤24 months), intermediate survival (Group 2, 24–48 months), and long survival (Group 3, 48–60 months). Differences among the three groups were evaluated using the Kruskal–Wallis test, with Benjamini–Hochberg false discovery rate (FDR) correction across biomarkers. Pairwise group comparisons were performed using Dunn’s post hoc test with FDR adjustment. Center lines indicate medians, boxes represent the interquartile range, whiskers extend to 1.5 times the interquartile range, and circles denote outliers. Pairwise significance is indicated as *p* < 0.05 (**), p < 0.01 (**), p < 0.001 (****), and not significant (ns).

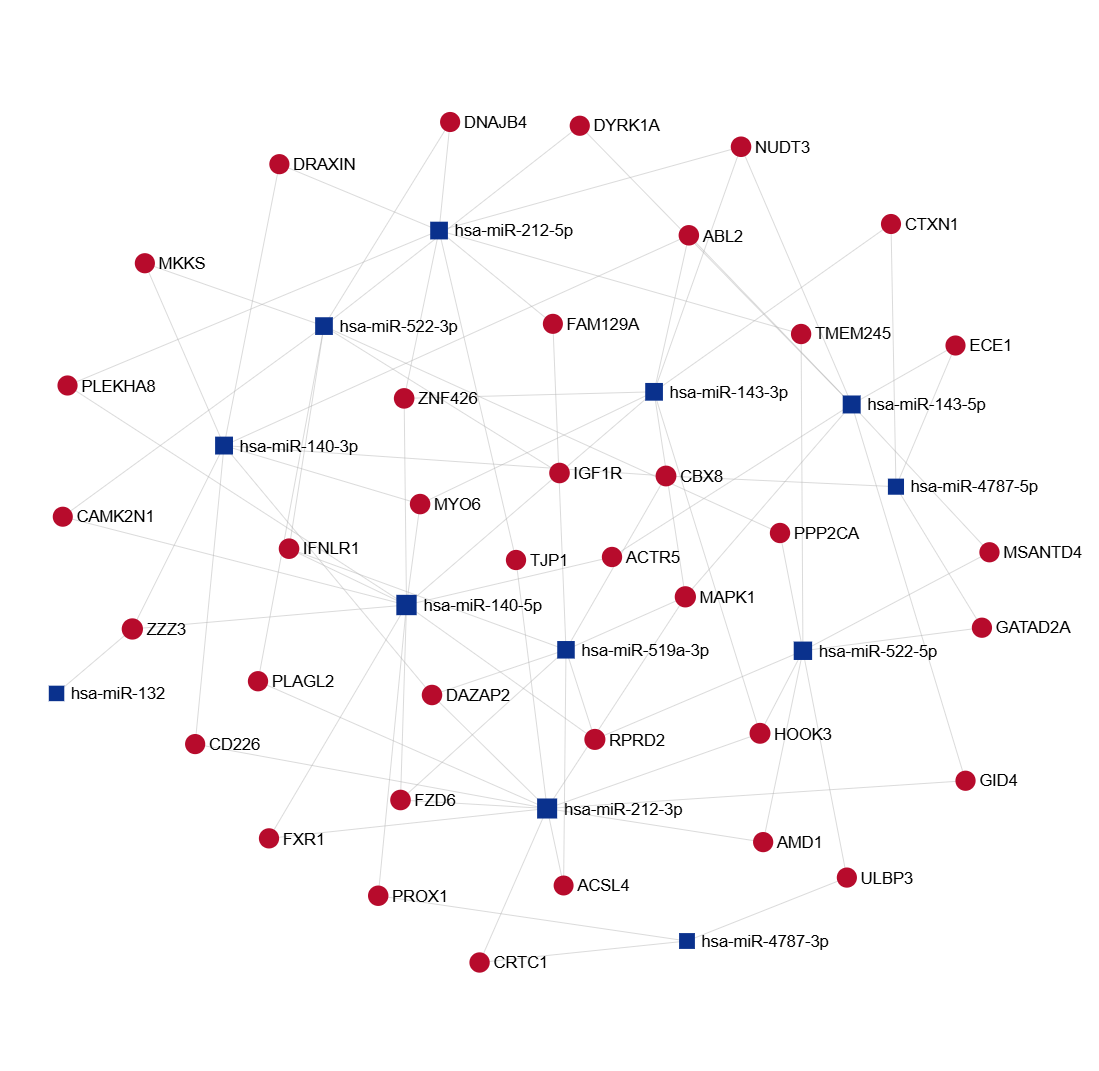
**Supplementary Figure S3. miRNA–target gene interaction network.** The seven machine learning-selected miRNAs (dark blue squares) are connected to their predicted target genes (dark red circles). For visualization purposes, the network was simplified to display the minimal set of mapped target genes while preserving the interactions among the selected miRNAs.

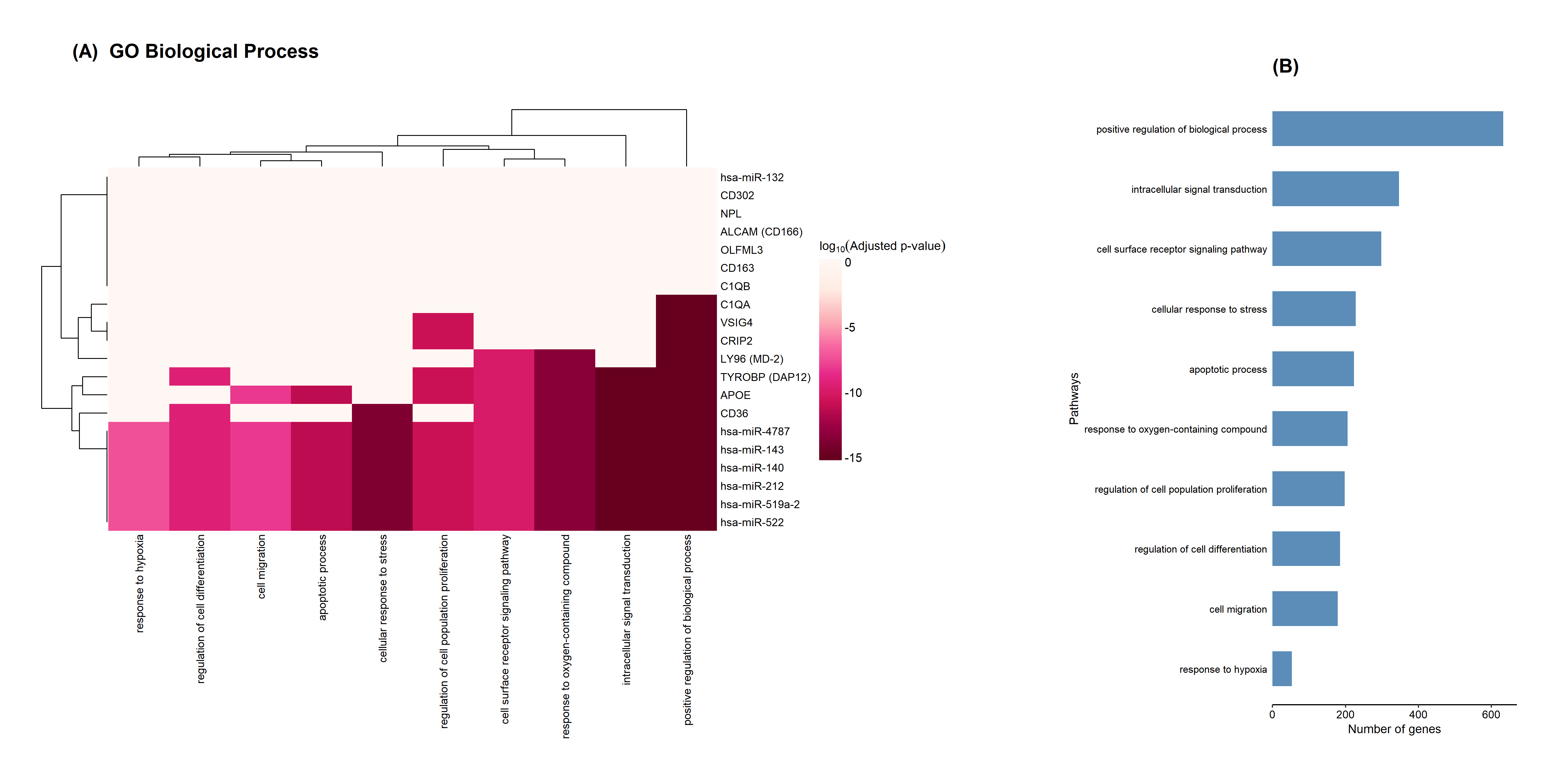
**Supplementary Fig. S4. GO Biological Process Landscape of Candidate Ovarian Cancer Biomarkers.** The heatmap (A) illustrates the association of candidate mRNA and miRNA biomarkers with the top 10 enriched GO biological processes. Colored cells indicate that a biomarker is associated with a given pathway, with color intensity representing the adjusted enrichment *p*-value (darker colors indicate greater statistical significance). Hierarchical clustering groups biomarkers and pathways according to their association patterns. The bar plot (B) shows the number of genes contributing to each enriched GO biological process.

**Supplementary Tables**

**Supplementary Table S1.** Comparison of representative recent studies on multi-omics integration and machine learning for ovarian cancer prognosis.

| **Study** | **Cohort** | **Omics layers** | **Preprocessing** | **Feature selection** | **Model** | **Performance** | **Ref.** |
| --- | --- | --- | --- | --- | --- | --- | --- |
| Zhang et al., 2016 | TCGA-OV (599) | mRNA + miRNA + methylation + CNV | Cox regression P < 0.05; pathway screening | PAM, hierarchical clustering, NMF | KM survival P values | 9 RNA-seq subtypes, 4 methylation subtypes | [57] |
| El-Manzalawy et al., 2018 | TCGA-OV (215) | RNA-seq + methylation + CNA | Complete-case intersection | Stage I LASSO; Stage II MRMR-mv | RF, XGBoost, logistic regression | Best multi-view AUC ≈ 0.70 | [14] |
| Hira et al., 2021 | TCGA-OV (292, 481) | RNA-seq + methylation + CNV | Sample intersection; zero-variance filtering | Variational autoencoder | VAE, MMD-VAE, Cox, SVM | MMD-VAE outperformed VAE | [66] |
| Wang et al., 2023 | TCGA-HGSOC (512) | RNA-seq + miRNA + methylation + CNV | Low-expression filtering; log transform | Survival-associated feature screening | MWINA clustering; KM/chi-square | OS chi-square P < 4.21 × 10⁻¹¹ | [67] |
| Wang et al., 2024 | TCGA-OV (325); GSE26712 | RNA-seq + miRNA + methylation + CNV | Removed features with >20% missing | RF + LASSO | MDCADON multi-modal DNN | TCGA accuracy 69.4% | [68] |
| Eldesouki et al., 2025 | TCGA-EOC (2,427) | RNA-seq + genomics + clinical + mutation | MICE imputation; IQR/Z-score filtering | Chi-square tests, feature ranking | Logistic regression, SVM, RF, GBC | GBC AUC = 0.81; RF AUC = 0.79 | [18] |
| Ma et al., 2025 | TCGA-OC (276); GEO | RNA-seq + miRNA + CNV + mutation | Omics matching; log/TPM normalization | MOVICS screening (top 1,500) | 10 multi-omics clustering algorithms | KM survival, C-index, calibration | [69] |
| Zhou et al., 2025 | TCGA-OV (267); GSE9891 | mRNA + lncRNA + miRNA + mutation | Removed incomplete samples; ComBat | MOVICS top 1,000 variable features | 10 MOVICS clustering algorithms | CS1/CS2 subtypes; 5-gene signature | [15] |
| Fu et al., 2018 | TCGA-OV | Somatic mutation + mRNA expression | Cancer-relevant gene filtering | Minimally adjusted Cox screening | Polygenic Cox models | 5-year OS AUROC = 0.80 | [19] |
| Bhardwaj & Van Steen, 2020 | TCGA-OV (230) | Gene expression + methylation | Sample matching; invalid survival removed | MSigDB knowledge-based gene sets | DAI, iDRW, netDx, SSL, SNF | Best DAI AUC = 0.89 | [70] |
| **This study** | **TCGA-OV (329 / 168)** | **mRNA + miRNA + methylation + CNV + RPPA** | **Within-fold MAD filter; median imputation** | **Composite Pearson + Spearman + MI** | **3-stage stacked weighted ensemble** | **r = 0.752; C-index 0.779; Cox C-index 0.686** | **—** |

AUC, area under the curve; AUROC, area under the receiver operating characteristic curve; CNA, copy number alteration; CNV, copy number variation; DAI, Data and Analytics Integrator; DNN, deep neural network; EOC, epithelial ovarian cancer; GBC, gradient boosting classifier; HGSOC, high-grade serous ovarian cancer; KM, Kaplan–Meier; LASSO, least absolute shrinkage and selection operator; MAD, median absolute deviation; MI, mutual information; MICE, multiple imputation by chained equations; MMD-VAE, maximum mean discrepancy variational autoencoder; MOVICS, Multi-Omics Integration and Visualization in Cancer Subtyping; MRMR-mv, min-redundancy max-relevance multi-view; NMF, non-negative matrix factorization; OS, overall survival; PAM, partitioning around medoids; RF, random forest; RPPA, reverse-phase protein array; SNF, similarity network fusion; SSL, semi-supervised learning; SVM, support vector machine; TPM, transcripts per million; VAE, variational autoencoder. Reference numbers correspond to the main-text reference list.

**Supplementary Table S2. Univariate Cox proportional hazards regression analysis of clinical and molecular features associated with overall survival.**

| Feature | Hazard Ratio (HR) | 95% CI | p-value |
| --- | --- | --- | --- |
| Age | 1.378 | 1.188–1.599 | 2.39 × 10⁻⁵ |
| NOTCH1 | 0.835 | 0.710–0.982 | 0.029 |
| miR-3928 | 0.864 | 0.752–0.992 | 0.038 |
| C5AR1 | 1.172 | 0.977–1.405 | 0.087 |
| Residual Disease | 0.914 | 0.817–1.023 | 0.118 |
| miR-4522 | 1.083 | 0.942–1.245 | 0.262 |
| NFKBP65PS536 | 1.071 | 0.923–1.244 | 0.364 |
| VSIG4 | 1.090 | 0.900–1.319 | 0.380 |
| AMBP | 0.956 | 0.808–1.131 | 0.601 |
| PRLR | 1.023 | 0.888–1.178 | 0.752 |
| SYK | 0.977 | 0.843–1.133 | 0.757 |
| miR-132 | 1.023 | 0.849–1.234 | 0.809 |
| miR-140 | 0.980 | 0.813–1.181 | 0.834 |
| ELA3A | 0.985 | 0.838–1.157 | 0.852 |
